# Differentiation of mild cognitive impairment using an entorhinal cortex-based test of VR navigation

**DOI:** 10.1101/495796

**Authors:** David Howett, Andrea Castegnaro, Katarzyna Krzywicka, Johanna Hagman, Richard Henson, Miguel Rio, John A King, Neil Burgess, Dennis Chan

## Abstract

The entorhinal cortex is one of the first regions to exhibit neurodegeneration in Alzheimer’s disease, and as such identification of entorhinal cortex dysfunction may aid detection of the disease in its earliest stages. Extensive evidence demonstrates that the entorhinal cortex is critically implicated in navigation underpinned by the firing of spatially modulated neurons. This study tested the hypothesis that entorhinal-dependent navigation is impaired in pre-dementia Alzheimer’s disease.

Forty-five patients with mild cognitive impairment (26 with CSF Alzheimer’s disease biomarker data: 12 biomarker-positive and 14 biomarker-negative) and 41 healthy control participants undertook an immersive virtual reality path integration test, as a measure of entorhinal-dependent navigation. Behavioural performance was correlated with MRI measures of entorhinal cortex volume, and the classification accuracy of the path integration task was compared with a battery of cognitive tests considered sensitive and specific for early Alzheimer’s Disease.

Biomarker-positive patients exhibited larger errors in the navigation task than biomarker-negative patients, whose performance did not significantly differ from controls participants. Path-integration errors were negatively correlated with the volumes of the total entorhinal cortex and of its posteromedial subdivision. The path integration task demonstrated higher diagnostic sensitivity and specificity for differentiating biomarker positive versus negative patients (area under the curve = 0.90) than was achieved by the best of the cognitive tests (area under the curve = 0.57).

This study demonstrates that an entorhinal cortex-based virtual reality navigation task can differentiate patients with mild cognitive impairment at low and high risk of developing dementia, with classification accuracy superior to reference cognitive tests considered to be highly sensitive to early Alzheimer’s disease. This study provides evidence that navigation tasks may aid early diagnosis of Alzheimer’s disease, and the basis of this in animal cellular and behavioural studies provides the opportunity to answer the unmet need for translatable outcome measures for comparing treatment effect across preclinical and clinical trial phases of future anti-Alzheimer’s drugs.

## Introduction

To date, all interventional trials aimed at slowing the progression of Alzheimer’s disease (AD) have failed. Two of the main contributors to this failure are: i) problems in identifying the initial stages of AD, such that interventional trials are applied too late in the disease process, and ii) the lack of translatable outcome measures for comparing treatment effects across preclinical testing in animal models of disease and clinical trials in patient populations (Mehta *et al*, 2017).

Detection of AD-related changes in entorhinal cortex (EC) function provides a potential solution to both of these problems. Degeneration of the EC is a critical initial stage of typical AD (Braak and Del Tredici, 2015), with 60% loss of layer II EC neurons observed by the time cognitive impairment is manifest (Gomez-Isla *et al*, 1997). Additionally, there is emerging evidence that the initial stages of AD may be associated with the trans-neuronal spread of pathological tau within the EC-hippocampal circuit (de Calignon *et al*, 2012; Ahmed *et al*, 2014), prior to neocortical infiltration. As such, tests sensitive to EC function might have added value in identifying the very earliest stages of AD, prior to hippocampal involvement.

Extensive evidence from animal studies indicates that the EC is involved in spatial navigation. *In vivo* single cell studies have identified EC neurons with spatially-modulated firing patterns, including grid cells (Hafting *et al*, 2005), head direction cells (Sargolini *et al*, 2006) and border cells (Solstad *et al*, 2008), with firing activity coupled to spatial behaviours (McNaughton *et al*, 2006). Together with hippocampal place cells (O’Keefe and Dostrovsky, 1971), these EC cells are considered to represent the neural basis of a cognitive map (O’Keefe and Nadel, 1978) that mediates spatial behaviours (Fyhn *et al*, 2007; Lester *et al*, 2017). Within the EC, the medial EC is considered to be particularly involved in navigation, given that up to 95% of mEC neurons may be grid cells (Diehl *et al*, 2017), in contrast to lateral EC neurons which exhibit little spatial selectivity. Evidence that the EC underpins navigation in other mammalian species is supported by the demonstration of EC grid cells in bats (Yartsev *et al*, 2011), monkeys (Killian *et al*, 2012) and humans (Jacobs *et al*, 2013).

Spatial tests, based on the cognitive map theory, have already shown that spatial processing is impaired in early AD. The Four Mountains Test (4MT), a hippocampal-dependent test of allocentric spatial memory (Hartley, 2007), differentiates patients with mild cognitive impairment (MCI) with and without CSF biomarkers of AD (Moodley *et al*, 2015) and is predictive of conversion from MCI to dementia (Wood *et al*, 2016). Crucially for detection of AD prior to symptom onset, performance on the 4MT correlates with dementia risk score in asymptomatic 40-59 year olds (Ritchie *et al*, 2018), while young adult *APOE-e4* carriers at increased risk of AD exhibit reduced grid-cell like representations and activation of the EC during an fMRI navigation task (Kunz *et al*, 2015). Finally, impaired route learning and way-finding is observed in asymptomatic individuals with positive amyloid-PET scans (Allison *et al*, 2016).

These previous studies provide the backdrop for the present study, which investigates EC-dependent navigation in MCI patients at risk of developing dementia. Navigation will be tested using a path integration (PI) task (Fig. 1A), which assesses the ability to keep track of, and return to, a previously visited location. Accurate PI requires the continuous integration of multisensory cues (visual, proprioceptive and vestibular) representing current position and heading direction, in reference to a fixed location (Etienne and Jeffery, 2004; McNaughton *et al*, 2006). While several other brain regions have been implicated in PI, including the hippocampus, prefrontal and retrosplenial cortices (Chrastil *et al*, 2015, 2017), converging evidence from numerous sources indicates that the EC plays a critical role in PI. Human imaging studies demonstrate the EC’s role in components of PI such as route planning (Maguire *et al*, 1998; Jacobs *et al*, 2010), computation of goal direction (Chadwick *et al*, 2015) and goal distance (Spiers and Maguire, 2007; Howard *et al*, 2014), while animal studies demonstrate that PI is critically dependent on mEC grid cell activity (Campbell *et al*, 2018; Tennant *et al*, 2018).

**Fig 1.**
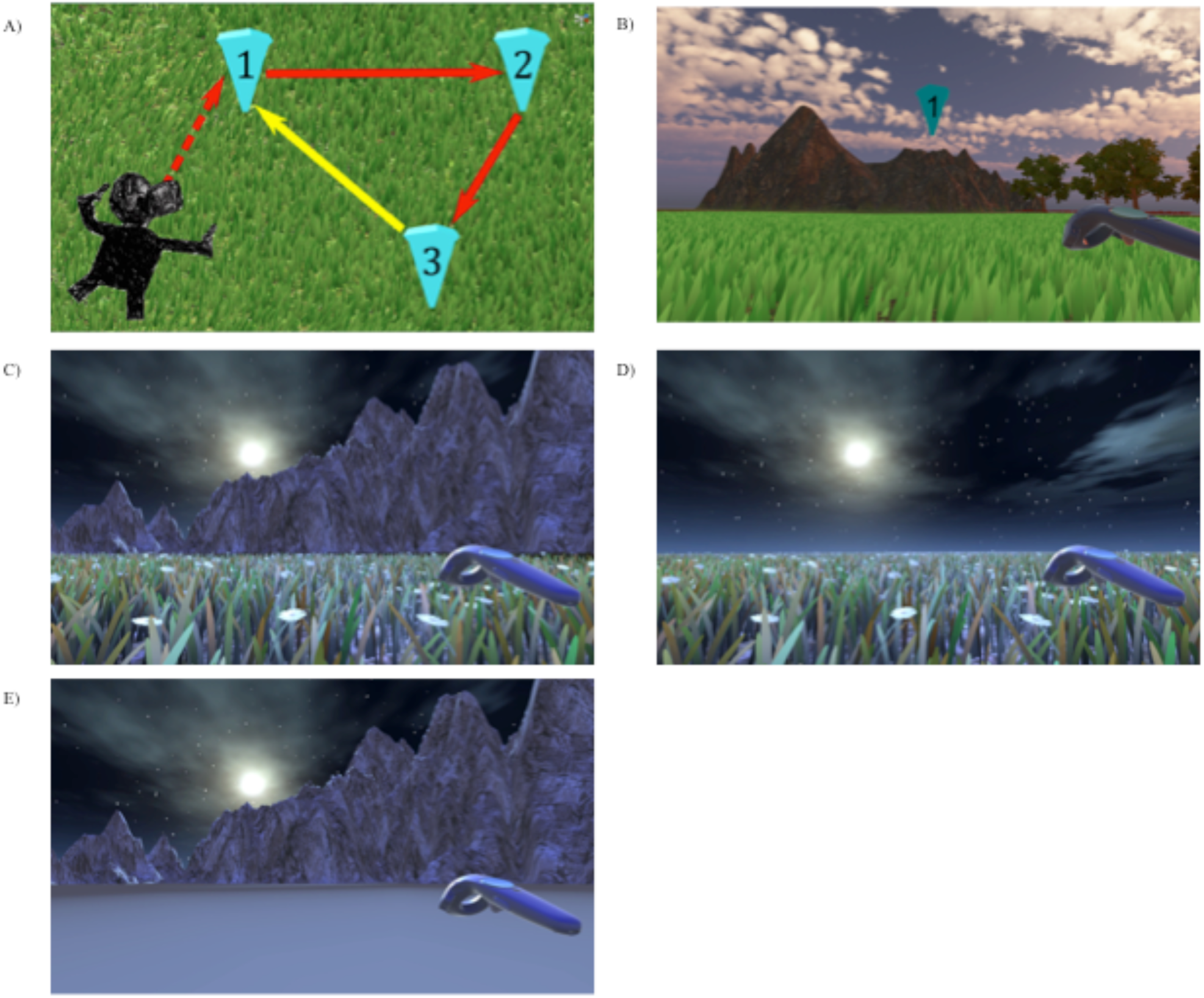
Path Integration Task. **A)** Illustration of the path integration task. Each numbered inverted blue cone is a location marker. Only one cone was visible at a time; upon reaching a blue cone it disappeared and the next one in the sequence appeared. Red arrows indicate the guided sequence along two sides of the triangle. The yellow arrow, the last side of the triangle, signifies the assessed return path, performed in the absence of any cones. **B**) Example environment from the head mounted display with textural and boundary cues present, with cone one and the controller shown. Texture and boundary cues are present in all trials when navigating between cones. **C-E**) Return conditions applied when attempting to return to the location of cone one only (yellow arrow (**A)**) and included no change (**C**), removal of environment boundaries (**D**) and removal of surface detail (**E)**.

In this study, PI will be tested using an immersive virtual reality (iVR) paradigm where participants navigate by real-world walking within simulated environments. Immersive VR has several theoretical and operational advantages over “desktop” VR tasks, which are typically performed seated and thus without locomotor or proprioceptive feedback, both of which are pivotal for grid cell function (Winter *et al*, 2015). First, the actual movement in iVR approximates real world navigation and thus has greater ecological validity than desktop VR. Second, there is evidence of differing neural processes underlying desktop and actual navigation, with desktop VR being associated with lower frequency hippocampal theta oscillations (Bohbot *et al*, 2017). This has negative implications for desktop VR as a valid proxy for real life navigation. Lastly, larger rotational (Klatzky *et al*, 1998) and distance errors (Distler *et al*, 1998; Sinai *et al*, 1999; Adamo *et al*, 2012) have been reported in desktop VR navigation when compared with tasks requiring active movement, possibly reflecting the absence of self-motion cues, leading in turn to reduced grid cell activation (Ólafsdóttir and Barry, 2015).

The primary objective of this study is to test the hypothesis that performance on an iVR PI task of EC function would differentiate MCI patients at increased risk of developing dementia. The secondary study objectives were to determine: i) whether manipulation of the environmental conditions would affect PI performance, ii) whether PI test performance correlates with EC volumes and iii) whether the PI task exhibits greater classification accuracy than current cognitive tests considered to have high diagnostic sensitivity and specificity for early AD.

## Materials and methods

### Participants

Patients with MCI (n=45) were recruited from the Cambridge University Hospitals NHS Trust Mild Cognitive Impairment and Memory Clinics. MCI was diagnosed by neurologists according to the Petersen criteria (Petersen, 2004), diagnosis of which requires; i) subjective cognitive complaint, ii) objective evidence of cognitive impairment, iii) preserved activities of daily living, iv) functional independence and v) absence of dementia. Objective cognitive decline was evaluated using the Addenbrooke’s Cognitive Examination – Revised (ACE-R, Mioshi *et al*, 2006) and a score of 0.5 on the Clinical Dementia Rating scale (CDR) (Morris, 1997). All patients underwent screening blood tests to exclude reversible causes of cognitive impairment. Exclusion criteria included the presence of a major medical or psychiatric disorder, epilepsy, a Hachinski Ischaemic Score > Four (Moroney *et al*, 1997), a history of alcohol excess or any visual or mobility impairment of such severity as to compromise ability to undertake the iVR test.

Twenty-six MCI patients underwent CSF biomarker studies (β-amyloid_1–42_, total tau, phosphorylated tau) as part of their clinical diagnostic workup. Biomarker studies were undertaken using ELISA assay kits (Innotest, Innogenetics, Ghent, Belgium) as outlined elsewhere (Shaw *et al*, 2009). Thresholds for negativity or positivity were set as CSF amyloid > 550pg/ml, CSF tau < 375pg/ml with a CSF tau: amyloid ratio of <0.8 (Mulder *et al*, 2010). MCI patients were stratified into biomarker-positive (MCI+, n=12) and biomarker-negative (MCI−, n=14) groups (Table 1). Researchers undertaking the VR tests were blinded to the CSF status of patients. The remaining 19 MCI patients did not undergo CSF studies as part of their clinical workup. Healthy control participants without a history of cognitive impairment (HCs, n=41, Table 1) were recruited from Join Dementia Research, an online repository of patients and volunteers interested in participating in dementia research.

**Table 1.**
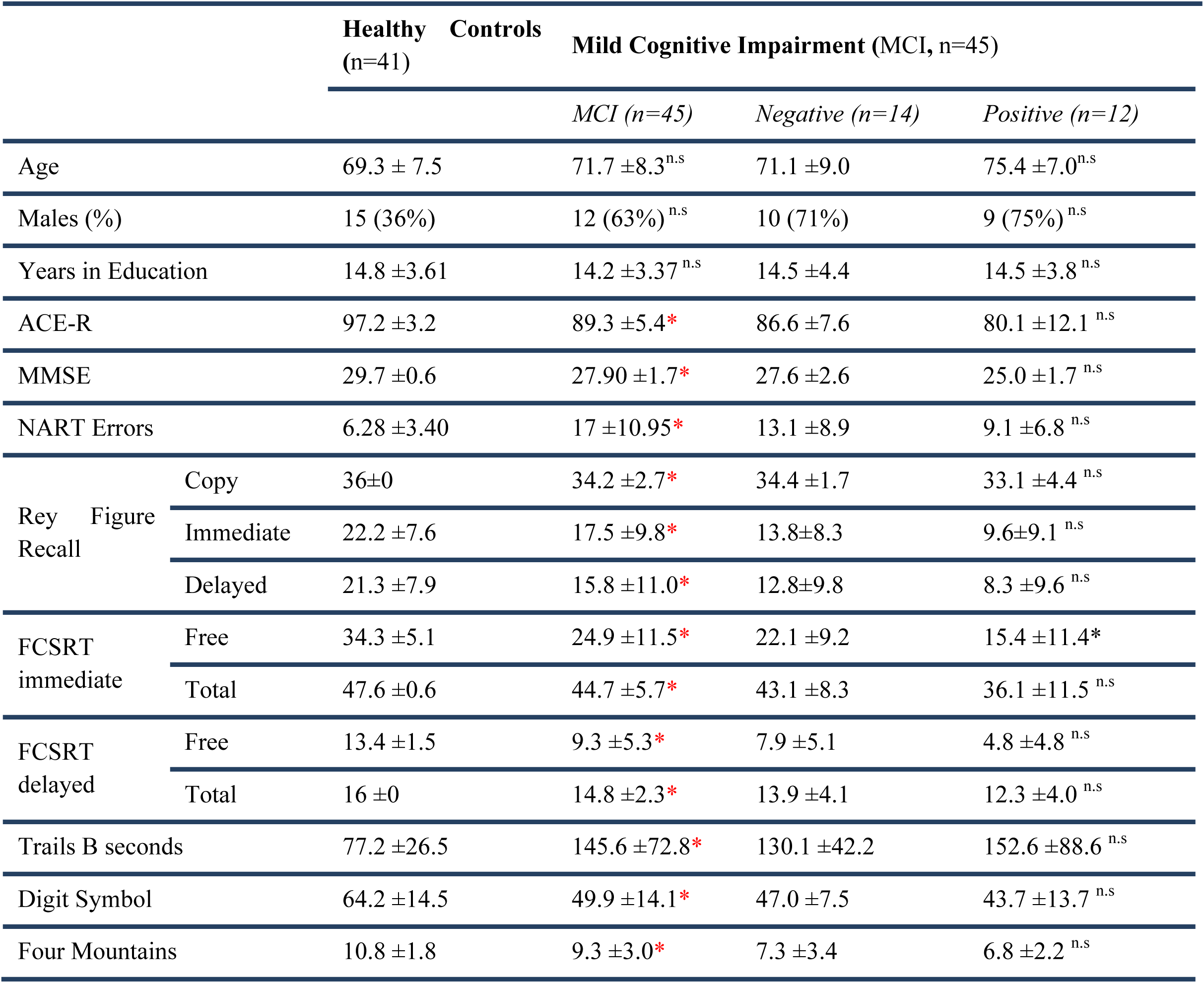
Demographics and neuropsychological test scores of across Controls and MCI patients. Table 1. Between group differences in neuropsychological test performance were assessed between HCs vs MCI as a whole, and MCI+ vs MCI−, scores indicate number of correct responses unless otherwise indicated. * = p<0.05, * = p<0.02 (Bonferroni-adjusted alpha); n.s = p>0.05). Abbreviations: 4MT – Four Mountains Test, ACE-R - Addenbrookes Cognitive Examination-Revised; MMSE - Mini-Mental State Examination; NART = National Adult Reading Test; FCSRT = Free & Cued Selective Reminding Test; Trails B = Trail Making Test B; Digit Symbol = Digit Symbol Substitution test.

The study was undertaken in line with the regulations outlined in the Declaration of Helsinki (WMA, 2013) and was approved by the NHS Cambridge South Research Ethics Committee (REC reference: 16/EE/0215).

### The iVR path integration task

The PI task was administered using the HTC Vive iVR kit, which uses external base stations to map out a 3.5×3.5m space within which participants walked during the VR task. If participants went beyond the tracked boundary by 30cm, an ‘out of border’ warning appeared in their sightline to encourage them to not walk any further. Researchers were also in the immediate proximity to ensure that participants did not venture beyond the test space.

The task was programmed in the Unity game engine and ran on Steam VR software, running on an MSI VR One backpack laptop.

The PI task was undertaken within virtual open arena environments with boundary cues projected to infinity (Fig. 1B). Three environments were used, each with unique surface details, boundary cues and lighting. The absence of local landmarks ensured EC-grid cell dependent strategies rather than striatal-mediated landmark-based navigation (Doeller *et al*, 2008).

No enclosure or local landmarks were present during the task in order to exclude any non-PI compensatory navigation strategies. A 1:1 correspondence between movement in the real and virtual worlds eliminated vestibular mismatch and minimised nausea and other tolerability issues.

Participants were asked to walk an ‘L’-shaped outward path to three different locations, each marked by inverted cones at head height numbered one, two and three (Fig. 1A and 1B). Only one cone was present at a time, each cone would disappear after the participant reached it and the next cone in sequence would appear. An auditory stimulus was presented with a cone’s appearance prompting participants toward the next cone’s location. Upon reaching cone three, a message projected onto the virtual scene asking participants to walk back to their remembered location of cone one (return path). When they reached their estimated location of cone one, participants pressed a trigger on a hand-held controller that logged their location and ended the trial.

Pre-trial practice sessions consist of 20 seconds of habituation to the iVR environment, during which participants were encouraged to explore the environment. Following habituation, participants performed five practice trials, where cone one was re-presented at the end of each trial in order to provide direct visual feedback to participants on the distance error between the remembered and actual locations of cone one.

The task consisted of nine trials conducted within each of the three environments, totalling 27 trials per participant. To examine the effects of environmental cues on PI, the environment was altered during the return path when participants were attempting to return to the remembered location of cone one. Three return conditions were used: (A) No environmental change (Fig. 1C), (B) removal of boundary cues (Fig. 1D, (C) removal of surface detail (Fig. 1E).

Each return condition was presented three times per environment, with return conditions presented pseudo-randomly in each environment in order to ensure participants were relying more on proprioceptive and self-motion cues rather than allothetic strategies.

Condition B was designed to increase dependence on self-motion cues and homing vector calculation by removing boundary cue information (Burgess *et al*, 2004), thereby placing a greater cognitive load on the PI network (Zhao and Warren, 2015). Condition C was designed to prevent feedback from surface motion during locomotion, thereby disrupting optic flow (Kearns *et al*, 2002) and increasing dependence on allocentric representations of space (Nardini *et al*, 2008). As such, return conditions B and C were considered analogous to “stress tests” for EC-network dependent navigation, with the prediction that a greater impairment in task performance would be observed during these conditions, compared with condition A.

Performance in the iVR PI task was assessed using three outcome measures. Absolute distance error, the primary outcome measure, was defined as the Euclidean distance between estimated and actual location of cone one, in line with previous research (Fig. 2, Chrastil *et al*, 2015; Mokrisova *et al*, 2016). Two secondary outcome measures were included to deconstruct absolute distance errors into its proportional angular and linear components. These measures additionally controlled for between-trial variance in triangle geometry owing to the pseudo-random generation of cone locations that could affect task difficulty, although variance is minimal in paths less than 10 metres long (Harris and Wolbers, 2012). Proportional angular errors reflect the accuracy of performed rotation at cone three toward the participant’s estimated location of cone one compared to the optimal rotation required to align with cone one (supplementary Fig. 1A). Proportional linear errors reflected the accuracy of distance estimation, with the Euclidean distance travelled between cone three and the participant’s estimated location of cone one compared to the distance between cone three and actual location of cone one (supplementary Fig. 1B).

**Fig 2:**
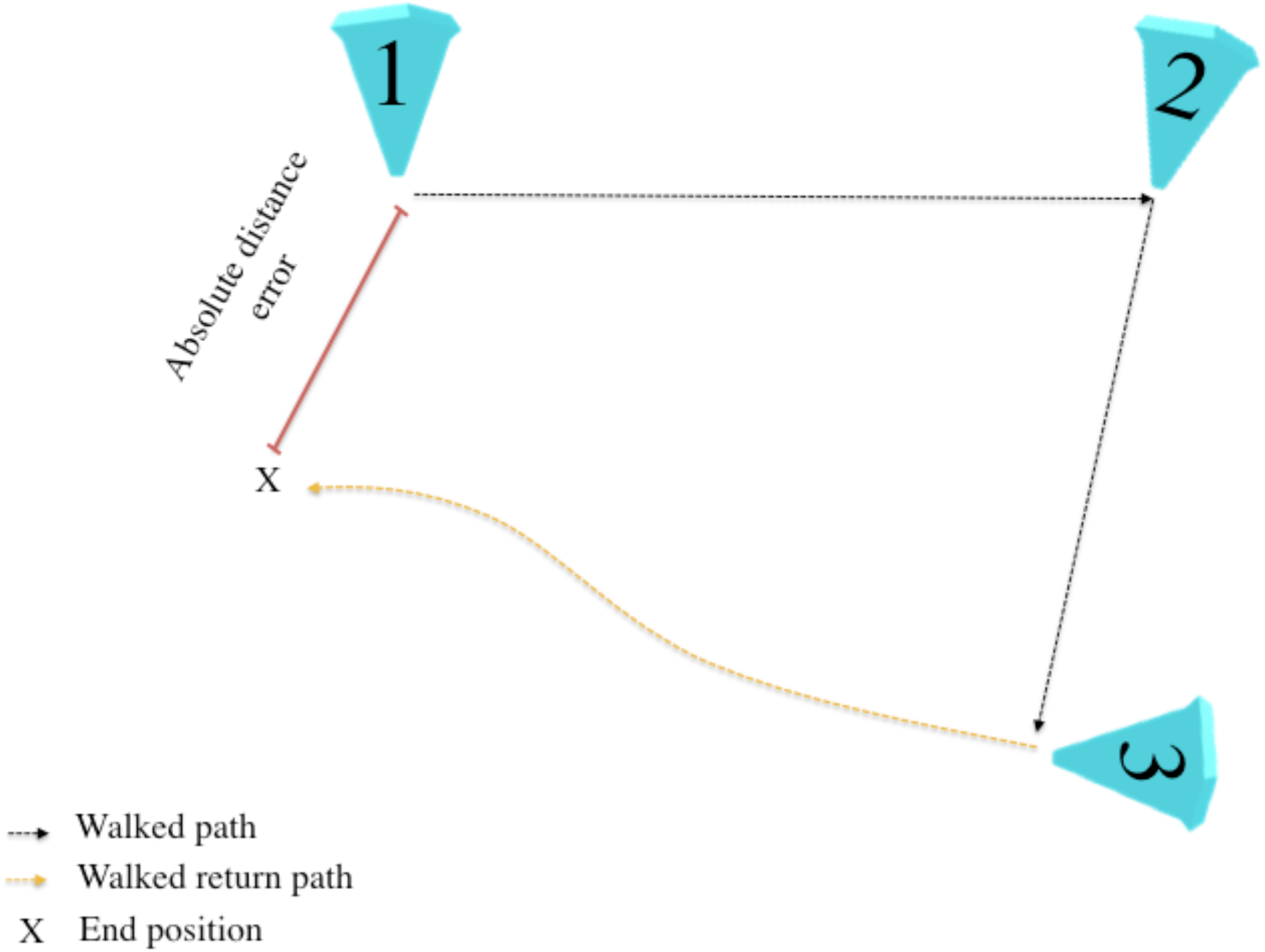
Primary measures of performance accuracy. Absolute distance error is defined as the Euclidean distance between the participant’s estimate of location one (goal) and the actual location of cone one.

### MRI Acquisition and analysis

37 HCs and 34 MCI patients (11 MCI+, 9 MCI−) underwent MRI scanning on 32 channel Siemens 3T Prisma scanners based either at the MRC Cognition and Brain Sciences Unit, Cambridge, or the Wolfson Brain Imaging Centre, Cambridge, with the same acquisition parameters used at the two scan sites. The scan protocol included whole brain 1×1×1mm T1-weighted MPRAGE (TA 5:12, TR 2300ms, TE 2.96ms) and high-resolution 0.4×0.4×2 mm T2-weighted scans through the hippocampal formation with scans aligned orthogonally to the long axis of the hippocampus (TA 8.11, TR 8020ms, TE 50ms).

As well as segmentation of the whole EC, additional segmentation of the anterolateral EC (alEC) and posteromedial EC (pmEC) subregions, representing the human homologues of the rodent lateral and medial EC respectively, was undertaken given their differing roles in spatial processing (Van Cauter *et al*, 2013; Knierim *et al*, 2014). While segmentation protocols for these subregions are available at 7T (Maass *et al*, 2015), no protocol is available for segmentation at 3T. Therefore, for this study, an in-house protocol was devised, which partially segmented alEC and pmEC using the three anterior-most and three posterior-most slices of the EC. Intermediate slices were not used for alEC and pmEC segmentation due to the overlap of the two subdivisions within this part of the EC. As such, this protocol prioritised specificity of segmentation over completeness (see supplementary methods). Manual segmentation was performed in ITK-SNAP (Yushkevich *et al*, 2006) (Fig. 3, detailed protocol in supplementary information). High inter- and intra-rater reliability was achieved for the manual segmentation protocol of the EC, alEC and pmEC, consistent with previous research (Berron *et al*, 2017; Olsen *et al*, 2017) - see supplementary table 1.

**Fig 3.**
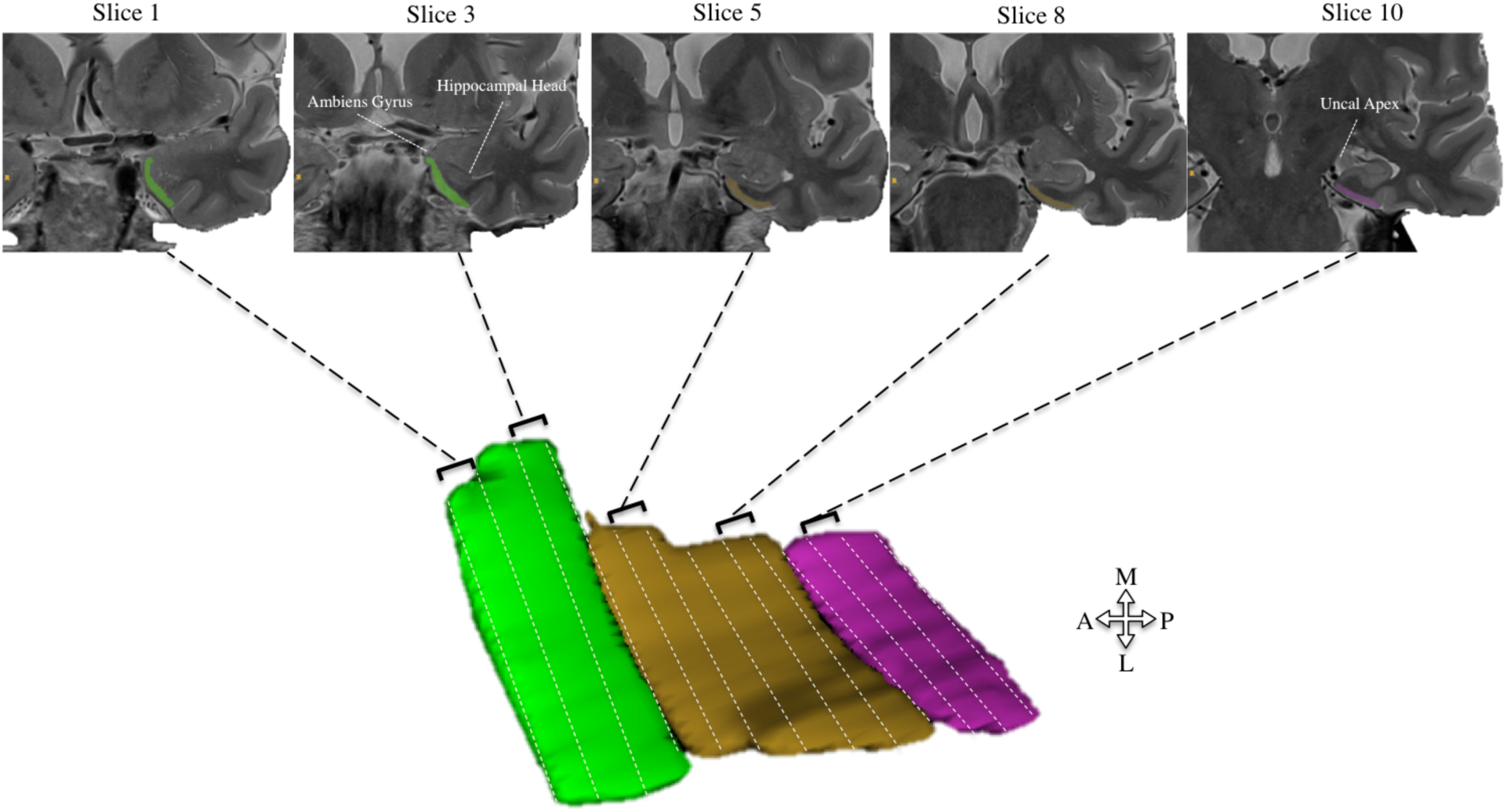
Illustration of volumetric segmentation of the anteriolateral, posterioromedial and total entorhinal cortex. Anterior-lateral (alEC, green) is segmented two slices anterior to the emergence of the hippocampal head (slice 3). Posteromedial (pmEC, pink) is segmented from one slice anterior to, and one slice posterior from, the uncal apex (slice 10). All intermediate slices between alEC and pmEC are segmented as EC (brown), total EC volume is produced by summing all 3 EC subdivision volumes. Arrow schematic indicates anatomical plane for 3D segmentation.

Given the involvement of the hippocampus and retrosplenial cortex (RSc) in PI (Worsley *et al*, 2001; Chrastil *et al*, 2015) these additional regions of interest (ROIs) were also segmented using Freesurfer 6.0 (Fischl *et al*, 2002; Iglesias *et al*, 2015). In the absence of an automated protocol for the segmentation of the RSc, the posterior cingulate cortex, which includes the RSc, was segmented as a proxy measure of the RSc.

All segmentations were manually inspected to exclude cysts, CSF and meninges; all volumetric measurements were averaged between hemispheres and normalised to intracranial volume.

### Comparator neuropsychological tests

To compare the ability of the iVR test to classify prodromal AD with that of reference neuropsychological tests considered to be highly sensitive to early AD. All participants were administered a battery of tests chosen for their effectiveness in predicting conversion from MCI to dementia (A-C), inclusion in the Preclinical Alzheimer’s Cognitive Composite (A and D) approved by the FDA for use as cognitive outcome measures in trials aimed at preclinical AD, or prior work indicating high sensitivity and specificity for prodromal AD (E). These tests are as follows (cognitive domains assessed in parentheses):

A. Free and Cued Selective Reminding Test (FCSRT, episodic memory – verbal, Buschke, 1984)
B. Rey figure recall (RFR, episodic memory – nonverbal, Osterrieth, 1944)
C. Trail Making Test B (TMT-B; executive function, attention, processing speed, Bowie and Harvey, 2006)
D. Digit Symbol test (DST, attention, processing speed, Ryan and Lopez, 2001)
E. 4 Mountains Test (4MT, allocentric spatial memory, Hartley *et al*, 2005)

All participants also underwent global cognitive testing with the Addenbrookes Cognitive Examination-Revised (ACE-R, Mioshi *et al*, 2006) and the National Adult Reading Test (NART, Nelson, 1982), as a measure of premorbid IQ.

### Statistical Analysis

Demographic differences between MCI+, MCI− and HCs were assessed using one-way ANOVA, or the Kruskal Wallis test where parametric assumptions were violated, whereas differences between HCs and total (combined) MCI were assessed using t-tests or non-parametric Mann-Whitney test.

Between-group performance in the PI task compared all MCIs against HCs, as well as MCI+ against MCI−. Linear mixed effect modelling (LME) was used to assess the effect of MCI status on absolute distance error, proportional angular error and proportional linear error. LMEs are the most suitable method for analysing clustered datasets (27 trials with one of three return conditions per trial per participant), with missing data (excluded due to travelling ‘out of border’, see Results), and unbalanced designs (Moen *et al*, 2016). Final model fixed effects included an interaction term between diagnosis and return condition, along with covariates of age, sex, years in education, ACE-R, NART and VR environment. Unique participant identifiers were used as the random intercept and VR environment as random coefficient, for further details see supplementary methods. Reported denominator degrees of freedom were computed using the conservative Satterthwaite approximation.

Between-group differences in ROI volumetry and cognitive performance across the neuropsychological test battery were investigated using one-way ANCOVA - rank ordered where parametric assumptions were violated (Conover and Iman, 1982) - covarying for age, sex and years in education. Separate linear regression models were used to assess absolute distance error (averaged per participant across all trials) and ROI volumetry, adjusting for age, sex, years in education and mean PI performance per participant group. Analyses were conducted across all participants and between MCI+ vs MCI-. Bonferroni correction was used to control for planned multiple comparisons. Residuals were visually inspected for violating linear assumptions, leverage and outliers.

The classification ability (MCI from HCs or MCI+ from MCI−) of the PI test (using per-participant averages of each PI outcome measure) was compared to reference cognitive tests. All linear classification models were adjusted for age, sex and years in education and used k-fold cross-validation (k=10) to control for over-fitting (Hawkins *et al*, 2003). Posterior probabilities following cross-validation were used to generate Area Under the Curve (AUC) of the receiver operating characteristic, as well as optimal sensitivity and specificity. Pointwise confidence intervals were generated following bootstrapping with 1000 replications.

All analysis was conducted in Matlab 2017b (Mathworks, https://uk.mathworks.com/).

## Data Availability

Anonymized data are available on request.

## Results

### Demographics and neuropsychological testing

No significant differences in age, gender, or years in education were observed between all MCI and HCs, or between MCI+ and MCI− (Table 1). Following Bonferroni correction with an adjusted α of 0.002, the MCI group as a whole exhibited significantly more errors in all neuropsychological tests compared to HCs (p<0.002), whereas no difference between MCI+ and MCI− survived multiple comparisons (p>0.002).

### Immersive VR path integration task

775 of the 2295 trials were excluded (33.77%) due to the ‘out of border’ boundary being reached during the return path, leaving 1520 viable trial remaining for analysis, with no between group difference in ‘out of border’ warnings were observed (p>0.05). All participants successfully completed the PI task with no reported nausea or tolerability issues.

### Absolute distance error

The MCI group as a whole exhibited significantly larger absolute distance errors than the HC group (t(1,107) = 3.24, p<0.01, Fig. 4A), with an estimated 57.33±17.87cm increase in absolute distance error compared to HCs. MCI+ patients exhibited significantly larger absolute distance errors compared MCI− patients (t(1,163)= 4.69, p<0.001, Fig. 4B), with an estimated increase of 97.56±20.34cm compared to MCI-. ACE-R score correlated with absolute distance errors across HCs and total MCI patients (t(1,85)= 2.89, p<0.01) and across MCI+ and MCI− groups (t(1,26)= 4.01, p<0.01), with lower ACE-R scores being associated with greater distance errors.

**Fig 4.**
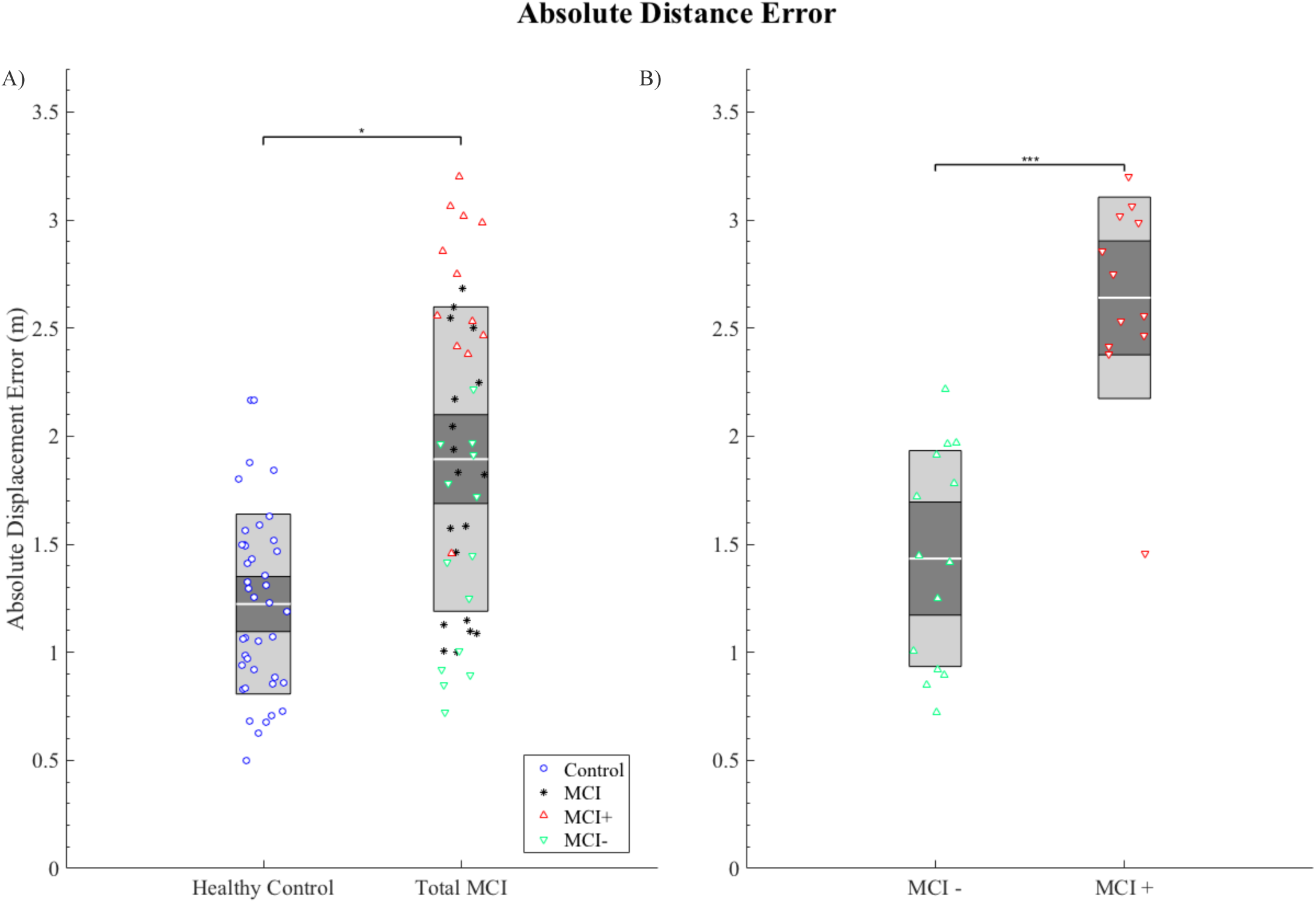
Graph summarising the between group differences in path integration performance. Absolute distance error (Euclidean distance) error in metres **(A-B)**. **A)** Group comparison between healthy controls and total MCI and B) between MCI− and MCI+. Each marker represents the mean performance across trials of each individual: blue circles = HCs; black asterisks = MCI without biomarkers; red triangles = MCI+; green inverted triangles = MCI−; Central gray line = mean; dark grey inner box= 95% confidence intervals; light grey outer box= 1 standard deviation. *p<0.05, ***p<0.001.

Similar main effects of diagnosis were observed in proportional linear errors between HCs and total MCI group (t(1,95)= 2.27, p<0.05), as well as between MCI+ and MCI− groups (t(1,87)= 3.09, p<0.001), but were not observed in proportional angular errors (p>0.05 across both groups; see supplementary results). No other fixed effect was associated with any of the outcome measures of the PI task.

### Effect of return condition on path integration

No main effects of return condition was observed on absolute distance error between HC and total MCI groups (F(2,1318)= 0.86, >0.05, Fig. 5) or between MCI+ and MCI− groups (F(2,384) = 0.56, p>0.05). Interaction terms between return condition and participant grouping were also included in the analyses to examine the differential influence of return conditions on PI performance for MCI+ (compared to MCI−) and MCI as a whole (compared to HCs). However, no interaction was observed between HC and total MCI groups (F(2,1325)= 0.83, >0.05) or between MCI+ and MCI− groups (F(2, 386) = 0.55, p>0.05).

**Fig 5.**
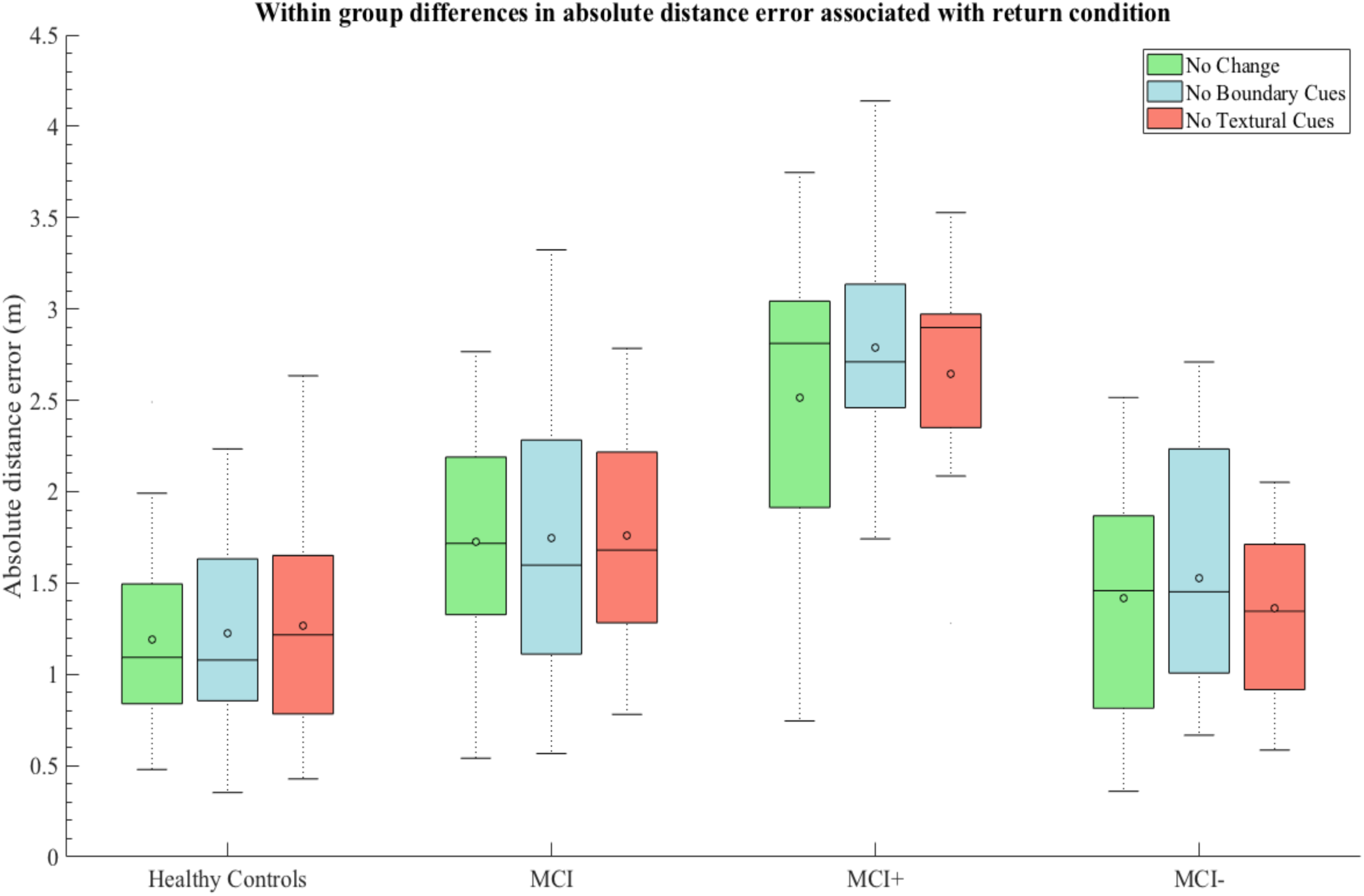
The effect of return condition within participant groups. The effect of return condition on absolute distance error averaged per participant in each group. Return conditions: Green = No environmental change; Blue = Removal of distal boundary cues; Red = Removal of surface detail. O = mean; black line = median.

For proportional angular errors, a trend toward an interaction between biomarker status (MCI+ vs MCI−) and return conditions B and C was observed (F(2,398) = 2.93, p<0.05, supplementary Fig. 4A), but this did not survive Bonferroni correction. No other main effect of return condition or interaction with diagnostic status was observed for proportional outcome measures (p>0.05, see supplementary results).

### Association between MRI volumetry and path integration performance

Group-level analyses adjusted for age, sex and years in education, revealed reduced ROI volumetry (PCC, hippocampus, EC, alEC and pmEC) in the total MCI group compared to HCs (p<0.05 for all ROIS), and in the MCI+ group compared to the MCI− group (p<0.05 for all ROIs). However, across HCs and total MCI, only hippocampal (F(1, 66)= 13.32, p<0.001), EC (F(1, 66)= 33.14, p<0.001), alEC (F(1, 66)= 21.87, p<0.001) and pmEC (F(1, 66)= 12.16, p<0.001) volumes survived the Bonferroni adjusted alpha of 0.005. See supplementary table 2 for full results.

Associations between ROIs and PI performance were assessed across total MCI and HC groups (Fig 6, purple line) and across MCI+ and MCI− groups (Fig. 6, grey line), adjusting for age, sex, years in education and average PI performance per participant group. Significant negative associations, surviving the Bonferroni adjusted alpha of 0.005, were observed across all participants between absolute distance errors and both total EC (F(1, 64)=9.60, p<0.005, Fig. 6A) and pmEC (F(1, 64)=9.73, p<0.005, Fig. 6C) volumes, each with an R^2^ of 0.38. Across HC and total MCI groups, associations between absolute distance error, alEC (p=0.04, Fig. 6B) and hippocampal (p=0.03, Fig. 6D) volumes did not survive Bonferroni correction. Similarly, for MCI+ and MCI− group comparisons, the association between hippocampal volume and absolute distance error (Fig. 6D, p=0.02) did not survive multiple comparison correction. No other ROI was associated with absolute distance error (p>0.05) across either participant grouping.

**Fig 6.**
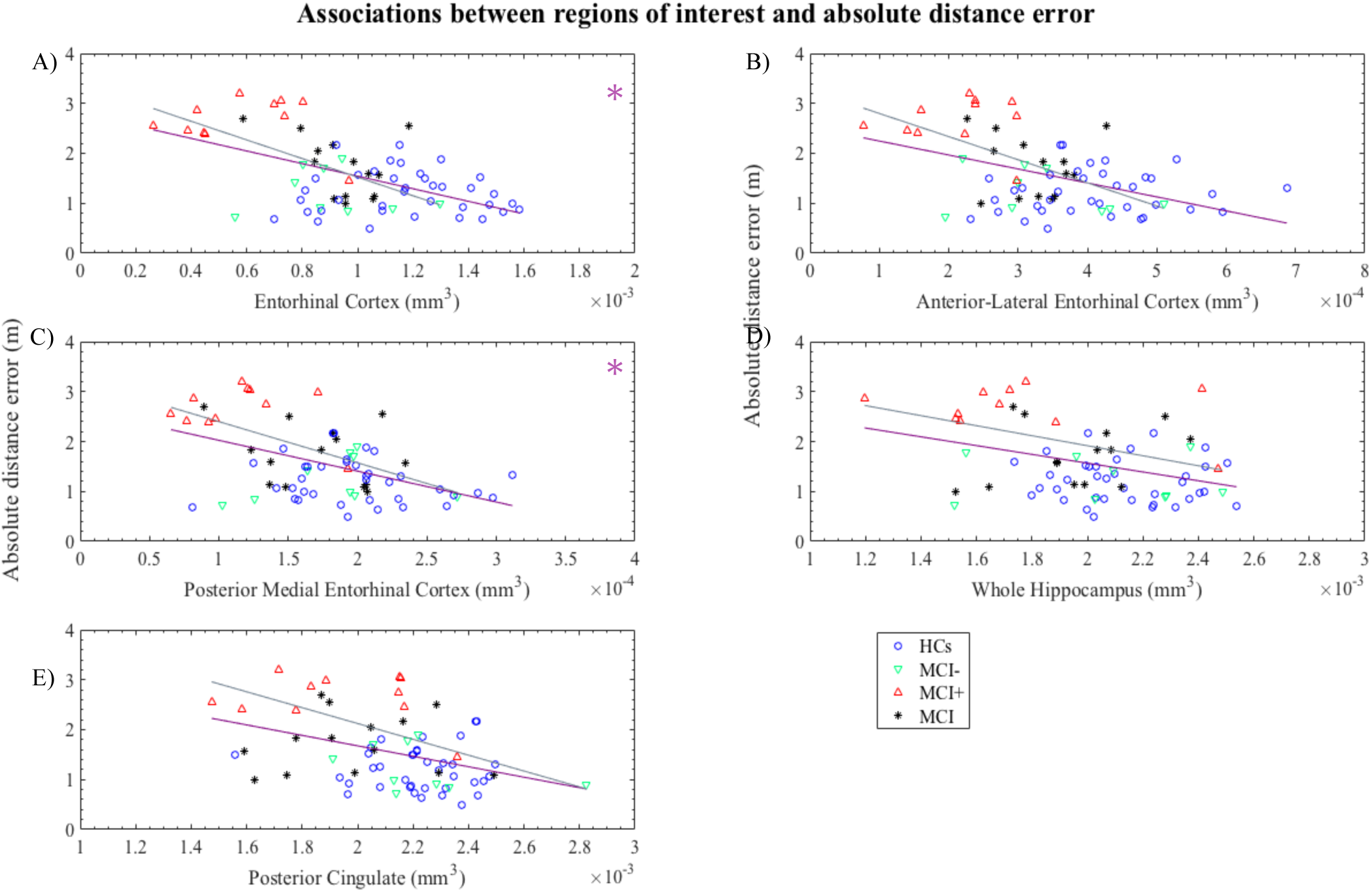
Scatterplots of path integration performance and region of interest (ROI) volumetry. The relationship absolute distance error and ROIs including; entorhinal cortex **(A)**, anterior-lateral entorhinal cortex **(B)**, posterior-medial entorhinal cortex **(C)**, whole hippocampus **(D)** and posterior cingulate cortex (E) was assessed. Least square lines are group specific: grey = across MCI+ and MCI−; purple= all participants), * in top-right corner of plot indicates p < 0.005 (Bonferroni adjusted α) across group as indicated by aforementioned colour.

### ROC curves and classification accuracy

Area under the curve (AUC), sensitivity and specificity were estimated using k-fold cross-validation (k=10), adjusted for age, sex and years in education. For the classification of total MCI patients from HCs, absolute distance error was associated with an AUC of 0.82 (Fig. 7A, 95% confidence intervals (CI) = 0.71-0.89), with an error ≥ 157cm yielding a sensitivity of 0.84 and specificity of 0.68. By comparison, the ACE-R was associated with an AUC= 0.86 (CI= 0.79-0.94)), TMTB (AUC= 0.79 (CI= 0.68-0.87)), 4MT (AUC= 0.73, (CI= 0.6-0.83)) and the delayed conditions of FCSRT (AUC= 0.73 (CI= 0.61-0.85)) and RFR (AUC= 0.72 (CI= 0.60-0.83)).

**Fig 7.**
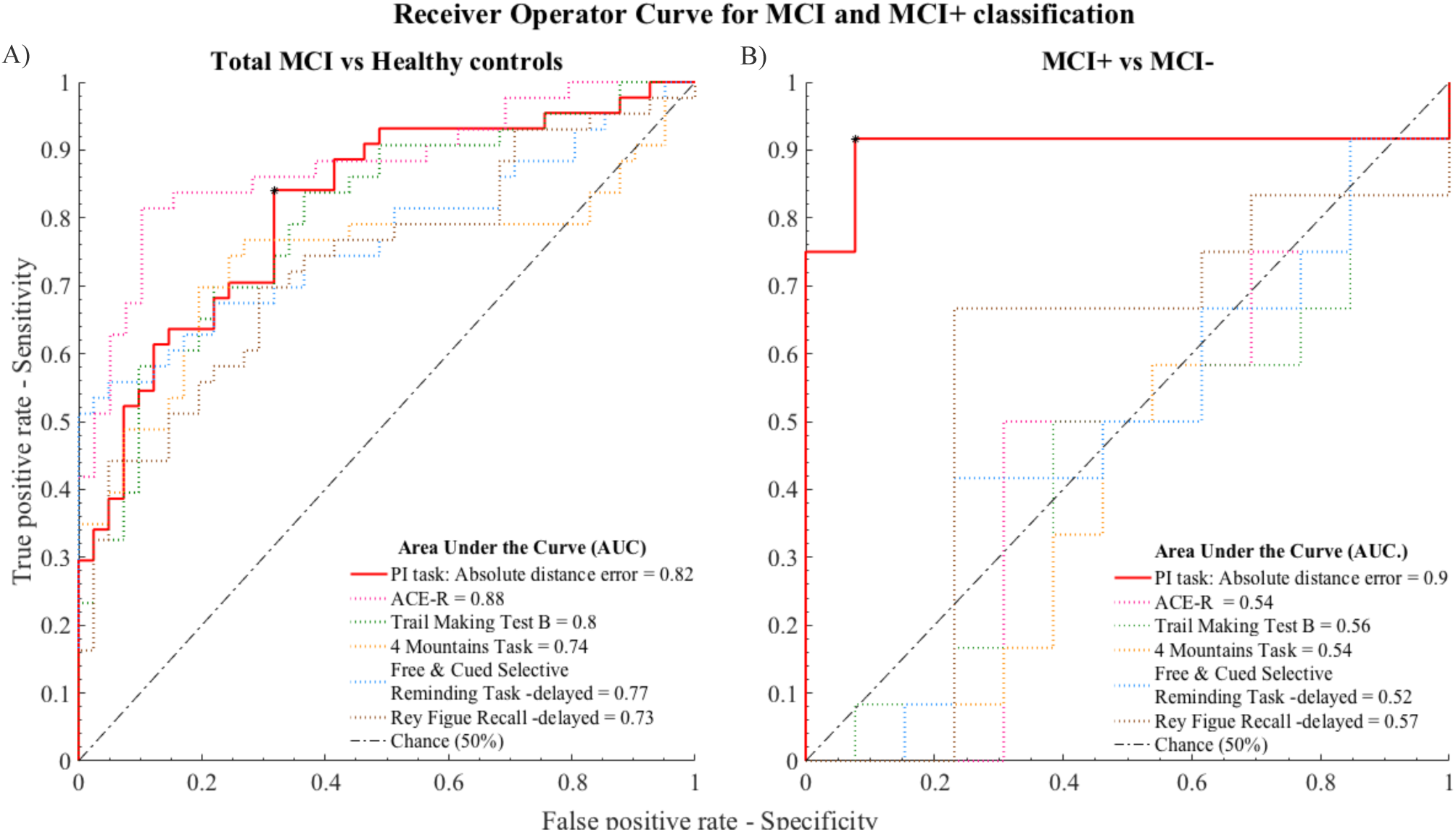
Receiver Operating Characteristic plot. Accuracy of path integration task performance for classifying **A)** total MCI from HCs and **B)** MCI+ from MCI-patients. PI performance is represented by absolute distance error (solid red line). Classification of reference cognitive tests is represented by dashed lines for comparison. Addenbrookes Cognitive Examination-Revised (grey), Trail Making Test B (green), 4 Mountains Test (yellow), Free and Cued Selective Reminding Test – delayed free recall (blue) and Rey Figure Recall – delayed recall (purple). Abbreviations: ACE-R - Addenbrookes Cognitive Examination-Revised. Asterix indicates optimal operating point for absolute distance error.

Classification accuracy of MCI+ from MCI− using absolute distance error was very high, with an AUC of 0.90 (Fig 7B, CI= 0.59-1), and errors ≥196cm yielding a sensitivity and specificity both of 0.92. This AUC was considerably higher than that of the comparator reference cognitive tests: ACE-R (AUC= 0.53 (CI= 0.24-0.73)), TMTB (AUC= 0.57 (CI= 0.22-0.69)), 4MT (AUC= 0.56, (CI= 0.22-0.72)) and the delayed conditions of FCSRT (AUC= 0.57 (CI= 0.22-0.68)) and RFR (AUC=0.55 (CI= 0.22-0.68)), indicating a markedly superior ability of the PI test to differentiate MCI+ from MCI−.

## Discussion

This study demonstrated that performance on a novel immersive virtual reality path integration paradigm, based on the central role of the entorhinal cortex in navigation, was impaired in MCI patients compared to healthy controls. In keeping with the study hypothesis that an EC-based navigation task can differentiate MCI patients at increased risk of developing dementia, we found that AD biomarker-positive patients drove the difference in navigation accuracy between MCI patients and controls. Consistent with the postulated role of the EC in navigation, and the specific role of the posteromedial EC subdivision (pmEC) in spatial processing, larger path integration performance errors were associated with smaller total EC and pmEC subdivision volumes across all participants. Finally, and of high relevance for potential diagnostic usage, PI performance differentiated MCI biomarker-positive patients, ie those with prodromal AD, from biomarker-negative patients with markedly higher sensitivity and specificity than a battery of “gold standard” cognitive tests used in clinical and research practice.

The navigational impairments observed in MCI patients is in line with previous navigation research (Hort *et al*, 2007; Laczó *et al*, 2014; Peter *et al*, 2018) and with the sparse literature on real-space PI in MCI and AD (Mokrisova *et al*, 2016). Significantly larger absolute distance errors were observed in MCI+ than in MCI−, with near-total separation of these two groups on this primary outcome measure, with the latter group exhibiting comparable performance to HCs. These data suggest that navigational deficits are relatively specific to AD and unrelated to deficits in other cognitive domains, such as attention or episodic memory, that might underlie the symptomatology of MCI− patients. Secondary outcome measures suggested that MCI+ patients are specifically impaired in distance estimation, as evidenced by reduced proportional linear errors, in line with previous research (Hort *et al*, 2007), and may relate to tau-related disruption of grid cell activity (Fu *et al*, 2017), given the role of grid cells in computing a distance metric of an environment (Bush *et al*, 2015) as part of path integration (McNaughton *et al*, 2006).

No group differences in performance errors were observed in response to the removal of boundary or surface detail cues. In the MCI+ group, a trend toward increased proportional angular errors in response to the removal of boundary (p=0.02) and textural (p=0.08) cues was observed, but this did not survive multiple comparison correction. Given that this effect did not reach corrected statistical significance, any inferences need to be made with caution. Nonetheless it is worth noting that this trend is consistent with previous research that reported heightened increased reliance on landmark cues (Kalová *et al*, 2005) and heightened rotational deficits in response to the disruption of optic flow (Kavcic *et al*, 2006; Mapstone *et al*, 2008).

Decreased volume was observed in the MCI group compared to controls in ROIs chosen for their role in path integration (total EC; including pm-EC and al-EC subdivisions, hippocampus and posterior cingulate cortex; a proxy measure of the RSc), although only significant differences in hippocampal, total EC, pmEC and alEC volumes survived correction for planned comparisons. In contrast to previous research (Dickerson and Wolk, 2013; Long *et al*, 2018) no difference in ROI volumetry across MCI+ and MCI− patients survived correction though this observation may be influenced by the sample sizes of these two groups. However, consistent with our hypothesis, both total EC and pmEC subdivision volumes were negatively associated with absolute distance errors across all participants, contrasting with the lack of association between this behavioural measure and alEC, hippocampal and posterior cingulate cortex volume. To our knowledge, this is the first demonstration that reduced pmEC volumes are associated with impaired path integration in humans, and is consistent with the analogous role of the rodent mEC in PI (McNaughton *et al*, 2006; Knierim *et al*, 2014). These findings reinforce previous work suggesting that the EC is critically involved in PI, and that PI in prodromal AD is primarily related to EC dysfunction rather than damage to the hippocampus or RSc, which are both involved in PI (Shrager *et al*, 2008; Kim *et al*, 2013) and affected in early AD (Braak and Braak, 1991).

PI performance differentiated the total MCI patient group from HCs with moderate classification accuracy (AUC 0.82), reflecting the large variance in performance within the former group. By comparison, PI performance was highly sensitive and specific for prodromal AD, classifying this group with an accuracy (AUC 0.90) that was markedly higher than that of reference cognitive tests of episodic memory, attention and processing speed widely used to diagnose prodromal AD and as outcome measures in clinical trial.

This work contributes to the growing body of evidence that spatial behavioural tests may have added value, above and beyond traditional cognitive tests, in detecting pre-dementia AD (Moodley et al. 2015, Allison et al (2016), Ritchie *et al*, 2018, Coughlan. *et al*, 2018). The use in this study of an EC-based navigation task potentially allows detection of AD in its very earliest stages, prior to hippocampal involvement, and builds on previous work showing alterations of EC activation during an fMRI navigation task in young adults at risk of AD due to *APOE-e4* genotype (Kunz *et al*, 2015).

This study has limitations. The sample size of both MCI+ and MCI− groups was relatively small, and these results therefore need to be considered initial findings that require replication in larger scale studies. Another limitation concerns the test space available with the commercial iVR hardware. The use of a larger space, which will be possible with next generation iVR, would likely result in the i) exclusion of fewer trials, ii) evaluation of proportional linear errors that is not skewed toward an undershoot, and iii) the compounding of vector computation errors (angular and linear estimates) that would likely culminate in larger between group performance differences. Finally, the lack of an anatomical mask for automated measurement of the retrosplenial cortex (RSC), necessitating the use in this study of a proxy anatomical measure (posterior cingulate cortex volume), limits analysis of the possible contribution of RSC dysfunction to the path integration impairment in prodromal AD.

In conclusion, this study demonstrates that performance on an EC-based iVR path integration task is sensitive and specific for prodromal Alzheimer’s disease, with greater classification accuracy than that of a battery of current “gold standard” cognitive tests. Given that this test is based on understanding of EC grid cell activity, these findings have implications not just for early diagnosis but also for translational AD research aimed at understanding mechanistic links between impaired cell activity and behaviour in AD. The task used in this study, combined with analogous navigation tasks in animal models of AD, would help address the need for outcome measures capable of comparing treatment effects across preclinical and clinical phases of future treatment trials aimed at delaying or preventing the onset of dementia.

## Supporting information

## Acknowledgements

The authors thank Daniel Bush for his comments on the manuscript.

## Funding

This work was supported by the Wellcome Trust Fellowship (202805/Z/16/Z), the European Research Council grant (ERC-2015-Adg, 694779) to NB. AC is sponsored by UCL’s Institute for Communications and Connected Systems. DH is supported by a UK Medical Research Council/Raymond and Beverly Sackler PhD studentship. RH is supported by MRC Programme Grant SUAG/010 RG91365. DC is funded by the Wellcome Trust, Isaac Newton Trust and the Cambridge NIHR Biomedical Research Centre.

## Competing Interests

Nothing declared.

## Abbreviations

4MT: Four Mountains Task
ACE-R: Addenbrookes Cognitive Examination – Revised
alEC: anterior-lateral Entorhinal Cortex
AUC: Area Under the Curve
CDR: Clinical Dementia Rating
EC: Entorhinal Cortex
DST: Digit Symbol Substitution Test
FCSRT: Free and Cued Selective Remind Test
HCs: Healthy Controls
iVR: immersive Virtual Reality
lEC: lateral Entorhinal Cortex
LME: Linear Mixed Effect Model
MCI: Mild Cognitive Impairment
MCI+: Mild Cognitive Impairment with CSF evidence of underlying Alzheimer’s Disease.
MCI−: Mild Cognitive Impairment without CSF evidence of underlying Alzheimer’s Disease.
mEC: medial Entorhinal Cortex
MMSE: Mini-Mental State Examination
NART: National Adult Reading Test
PCC: Posterior Cingulate Cortex
PI: Path Integration
pmEC: posterior-medial Entorhinal Cortex
RSc: RetroSplenial cortex
RFR: Rey-Osterrieth Figure Recall Test
ROC: Receiver Operating Characteristic
ROI: Region of Interest
TMTB: Trail Making Test B

