## Supplementary material for "Differentiation of mild cognitive impairment using an entorhinal cortex-based test of VR navigation"

### Methods

#### Secondary outcome measures

Relative measures of performance are included for two reasons: i) to deconstruct the primary PI outcome measure (absolute distance error) – into its angular and linear components, and ii) to control for the pseudo random generation of cone locations that creates between-trial variance. Errors are therefore relative where proportional values <1 reflects underestimates of rotation or distance and values >1 reflect overestimates of rotation or distance.

*Proportional Angular Error*

Proportional angular errors represent the relative degree of rotational inaccuracies performed at cone 3 toward cone. Errors are calculated as the ratio of rotation performed at cone 3 toward estimated location of cone 1 over the amount of rotation required at cone 3 for an optimal return to cone 1 (supplementary Fig. 1A).

*Proportional Linear Error*

Proportional linear errors reflect discrepancies in distance estimation as calculated by the ratio of distance between cone 3 and estimated location of cone 1 over the actual distance between cone 3 and cone 1 (supplementary Fig. 1B).

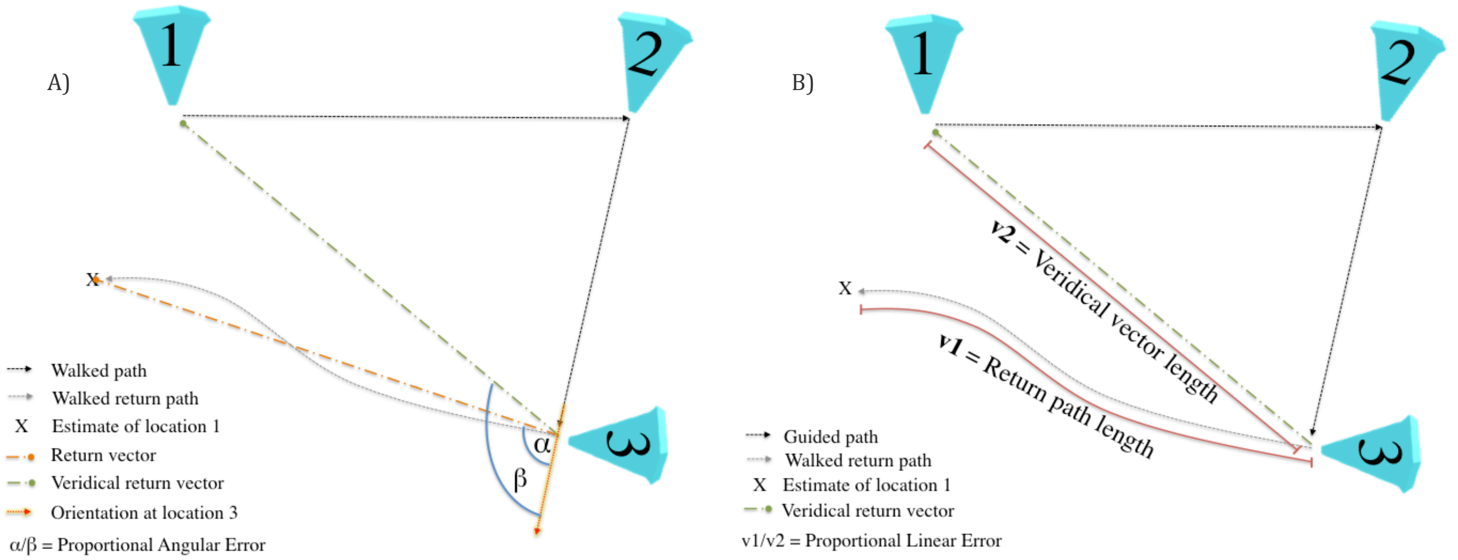


**Supplementary Fig 1. Illustrations of secondary outcome measures for the path integration task. A)** Proportional angular error is a measure of rotation accuracy, represents the ratio of performed rotation toward the estimated location of cone 1 at cone 3 (α) divided by the degree of rotation required toward cone 1's actual location (β). **B)** Proportional linear error represents the length of the performed return

#### Statistics – use of multilevel modelling.

With clustered data it is common practice to analyse performance by either i) pooling all trials together, thereby neglecting between-participant variance or ii) average observations for each participant, thereby neglecting within-participant variance. Both approaches may lead to misleading inferences and statements about statistical precision (Moen *et al.*, 2016).

The decision to use linear mixed effects modelling (LME) was supported by comparing model fit between empty linear and empty multilevel models using Likelihood Ratio testing. Separate models were used to assess PI performance in MCI+ vs MCI- vs HCs, and HCs vs pooled MCI, for absolute linear error, proportional angular error and proportional linear error. The final LME used was:

$$DV_{ij} = \beta_0 + \beta_1 MCI_j + \beta_2 Cond_{ij} + \beta_3 MCI_j * Cond_{ij} + \beta_4 Age_j + \beta_5 Sex_j + \beta_6 Edu_j + \beta_7 ACER_j + \beta_8 NART_j + \beta_9 Env_{ij} + U_{0j} + U_{1j} Env_{ij} + e_{ij}$$

where $DV_{ij}$ is the dependent variable (e.g. absolute distance error, separate models for each outcome measure) in trial *i (1...27)* of participant *j (1..86)*. $\beta_0$ is the population mean, $\beta_1 MCI_j$ is the diagnostic status of participant *j* (HC vs MCI or MCI+ vs MCI- vs HC), $\beta_2 Cond_{ij}$ is the return condition (no change, no boundary cues, no textural cues) of trial *i* for participant *j.* An interaction term between MCI status and return condition ($\beta_3$) is also included. There were additional fixed effects of participant's age ($\beta_4$), sex ($\beta_5$), years in education ($\beta_6$), ACE-R ($\beta_7$) and NART ($\beta_8$) scores. $U_{0j}$ is the random intercept for each participant, while the three different VR environments were modelled as a fixed effect ($\beta_9$), as well as random slopes that depended on participant ($U_{1j}$). The term $e_{ij}$ is the trial-level error for each trial *i* of participant *j*.

Visual inspection of residual plots did not reveal any significant deviations from homoscedasticity or normality.

Final LME models were informed by a mixture of *a priori* hypotheses and covariates, where appropriate they were refined using likelihood ratio testing for goodness of fit, whereas intraclass correlation coefficient was used to estimate the amount of variance explained by random effects.

#### Manual segmentation protocol

For this study an additional in-house protocol was used to segment the anterior-lateral (alEC) and posterior-medial (pmEC) EC subfields, which are analogous to the lateral and medial EC of rodents that have complementary roles in object and space processing respectively (Van Cauter *et al.*, 2013). The alEC and pmEC volumes are derived from the anatomical masks for application at 7T and developed to investigate the functional connectivity of this region (Maass *et al.*, 2015). However, given the progressive boundary between these functionally distinct subregions, we adopted to segment only the terminal most slices utilizing robust anatomical landmarks (Berron *et al.*, 2017) in order to avoid subfield mislabeling.

The EC, alEC and pmEC were manually segmented on coronal slices of high resolution T2-weighted 3T MRI scans of all participants, where MRI data was available, segmentation of EC subfields is summarized in Fig. 2 (main text). The alEC was segmented on the three anterior-most slices, whereas the pmEC was segmented on the three posterior-most terminal slices of the EC. The segmentation of alEC began two slices anterior to appearance of the hippocampal head and subiculum extending to the first slice of the subiculum. Whereas, pmEC segmentation extends one slice anterior from, to one slice posterior to the emergence of the incisura temporalis and the separation of the uncus from the medial temporal lobe.

#### Results

##### Reliability of manual segmentation

Intra-rater reliability was assessed for both raters by re-segmenting 5 randomly selected scans following a delay from initial segmentation (DH: 3 months and JH: 1 month). Inter-rater reliability was assessed for both raters by segmenting 5 randomly selected images from the other rater. Intra- and inter-rater reliability was assessed using spatial overlap as measured using Convert3D's Dice similarity coefficient (Yushkevich *et al.*, 2006). Whereas volumetric consistency was assessed using intraclass correlation coefficient (ICC) for intra- (ICC (3,k) -consistency) and inter-rater (ICC(2,k) -agreement) reliability (McGraw and Wong, 1996). Reliability metrics are presented in supplementary table 2, where intra-rater ICC represents the average of both raters.

**Supplementary table 1:** Reliability Measurements

| | | Entorhinal Cortex | | Anterior-lateral EC | | Posterior-medial EC | |
|---|---|---|---|---|---|---|---|
| | | LHS | RHS | LHS | RHS | LHS | RHS |
| Intra-rater reliability | *ICC | 0.93 | 0.82 | 0.88 | 0.89 | 0.85 | 0.77 |
| | Dice | 0.87 | 0.87 | 0.89 | 0.82 | 0.88 | 0.85 |
| Inter-rater reliability | ICC | 0.98 | 0.94 | 0.92 | 0.98 | 0.91 | 0.70 |
| | Dice | 0.70 | 0.71 | 0.75 | 0.73 | 0.71 | 0.69 |

Dice similarity coefficient was computed for both intra- and inter-rater reliability. ICC(3k) and ICC(2k) was used for intra-rater and inter-rater reliability, respectively. Abbreviations: EC, entorhinal cortex; ICC intraclass coefficient; *ICC – averaged across both raters.

#### Proportional Angular Error

No significant difference in proportional angular error was observed between either total MCI and HCs (T(1,128) = 1.79 p>0.05, supplementary Fig. 3A) or between MCI+ and MCI- (T(1,136) = 1.06, p>0.05, supplementary Fig. 3B). Although between total MCI and HCs, the fixed effect of MCI exhibited a non-significant trend toward a positive association with proportional angular error (p=0.07).

*Classification of proportional angular error*

Proportional angular errors exhibited an AUC of 0.77 (95% CI 0.65-0.86), with an error ≥ 1.12 yielding a sensitivity of 0.75 and specificity of 0.73.

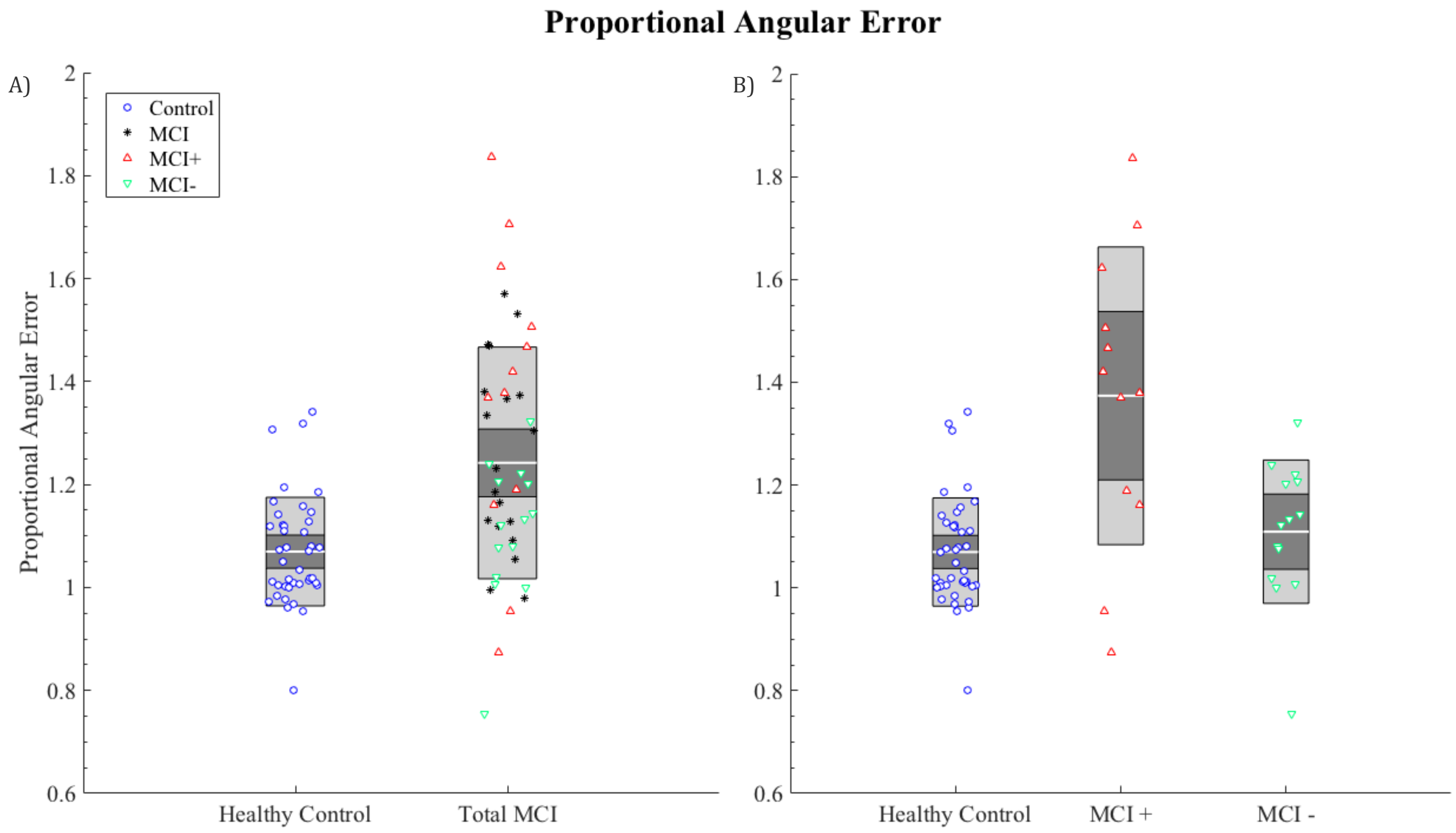


**Supplementary Fig 2. Graph summarising between group proportional angular error.** Performance was evaluated between **A)** total MCI and healthy controls and **B)** healthy controls, MCI+ and MCI-. Each marker represents the mean performance across trials of each participant: blue circles = HCs; black asterisks = MCI without biomarkers; red triangles = MCI+; green inverted triangles = MCI-. Central gray line = mean; dark grey inner box= 95% confidence intervals; light grey outer box= 1 standard deviation

#### Proportional Linear Error

Significant differences were observed between MCI and HCs ($T(1,95)= 2.27$, $p<0.05$, supplementary Fig. 2A), as well as between MCI+ and MCI- ($T(1,87)=3.09$, $p<0.01$, supplementary Fig. 2B). Compared to HCs, total MCI participants exhibited a decreased proportional linear error of 0.12±0.05, whereas MCI+ patients exhibited a decrease of 0.23±0.07 compared to MCI-.

##### Classification of proportional linear error

Proportional linear errors exhibited an AUC of 0.71 (95% CI 0.61-0.84), where errors ≥ 0.79 yielded a sensitivity of 0.66 and specificity of 0.78.

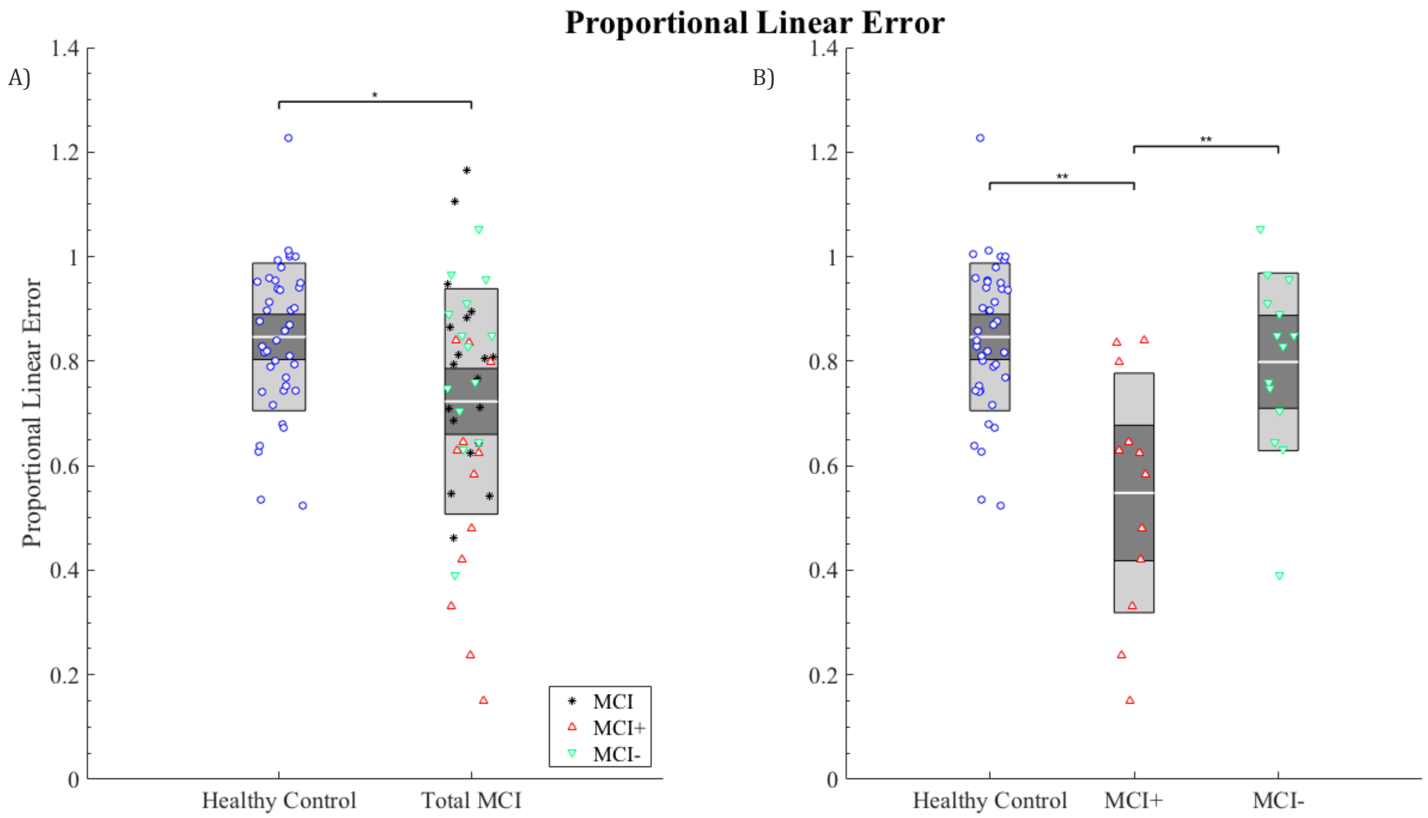


**Figure 3. Graph summarising between group differences in proportional linear error.** Proportional linear between pooled MCI and healthy controls **(A)** and healthy controls, MCI+ and MCI- **(B)**. Each marker represents the mean performance across trials of each participant: blue circles = HCs; black asterisks = MCI without biomarkers; red triangles = MCI+; green inverted triangles = MCI-; Central gray line = mean; dark grey inner box= 95% confidence intervals; light grey outer box= 1 standard deviation

#### Effect of return condition on secondary outcome measures

No significant main effect of return condition on proportional angular error was observed between HCs and total MCIs ($F(2,1317)= 1.37$ $p>0.05$) as well as across MCI+ and MCI- ($F(2,395)= 0.13$, $p>0.05$, supplementary Fig. 4A). However, a trend toward an interaction between biomarker status and return condition was observed on proportional angular errors ($F(2,398) = 2.93$, $p<0.05$, supplementary Fig. 4B), however this did not survive multiple comparison. No interaction between MCI status and return condition was observed across MCI and HCs on proportional angular errors ($F(2,1317) = 1.37$, $p>0.05$).

No significant main effect of return condition on proportional linear error was observed across HCs and all MCI patients ($F(2,1310) = 0.96$, $p>0.05$) or MCI+ and MCI– ($F(2,381) = 0.37$, $p>0.05$, supplementary Fig. 4A). No significant interaction was observed between MCI status and return condition on proportional linear error ($F(2,1317) = 0.38$, $p>0.05$), nor between biomarker status (MCI+ and MCI-) and return condition ($F(2,382) = 0.35$, $p>0.05$, supplementary Fig. 4B).

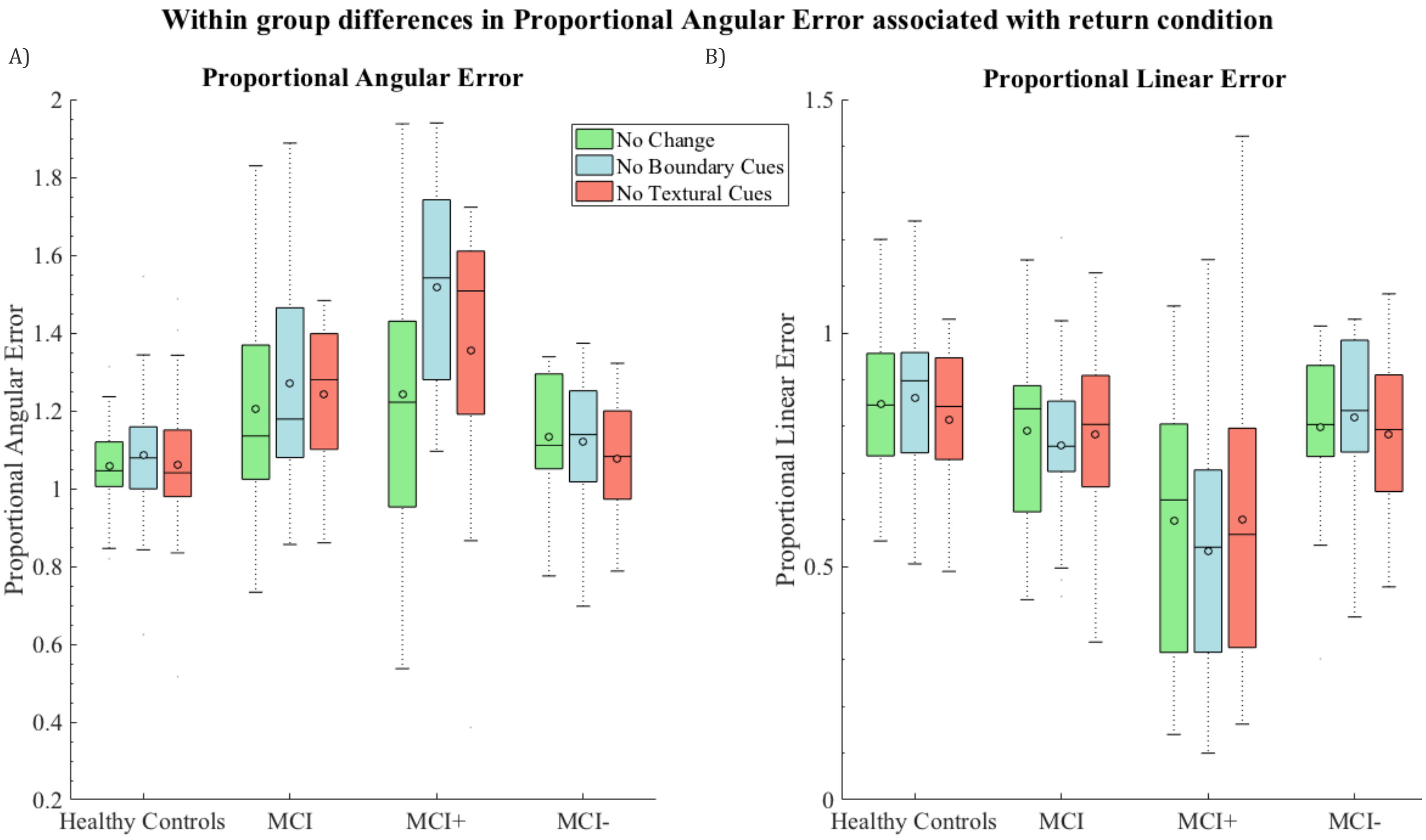


**Supplementary Fig 4. The effect of return condition between participant groups.** The effect of return condition on **A)** proportional angular error and **B)** proportional linear error averaged per participant in each group. Return conditions: Green = No environmental change; Blue = Removal of distal boundary cues; Red = Removal of textural cues. O = mean; black line = median.

**Table 2. Between Group differences in region of interest volumetry.**

| | *HC vs all MCI* | *MCI+ vs MCI-* |
|---|---|---|
| Posterior cingulate cortex | $F(1, 66) = 11.20$, $p<0.01$ | $F(1, 15) = 5.83$, $p<0.05$ |
| Hippocampus | $F(1, 66) = 13.32$, $p<0.001$ | $F(1, 15) = 6.02$, $p<0.05$ |
| Entorhinal cortex | $F(1, 66) = 33.14$, $p<0.001$ | $F(1, 15) = 14.51$, $p<0.01$ |
| Anterior-lateral entorhinal cortex | $F(1, 66) = 21.87$, $p<0.001$ | $F(1, 15) = 11.60$, $p<0.01$ |
| Posterior-medial entorhinal cortex | $F(1, 66) = 12.16$, $p<0.001$ | $F(1, 15) = 11.91$, $p<0.01$ |

ANCOVAs were ran across the two participant groups with the region of interest as the dependent variable, patient diagnostic status as the predictor variable with age, sex and years in education covariates. Bonferroni adjusted $\alpha = 0.005$.

Distribution of estimated goal location (cone 1)

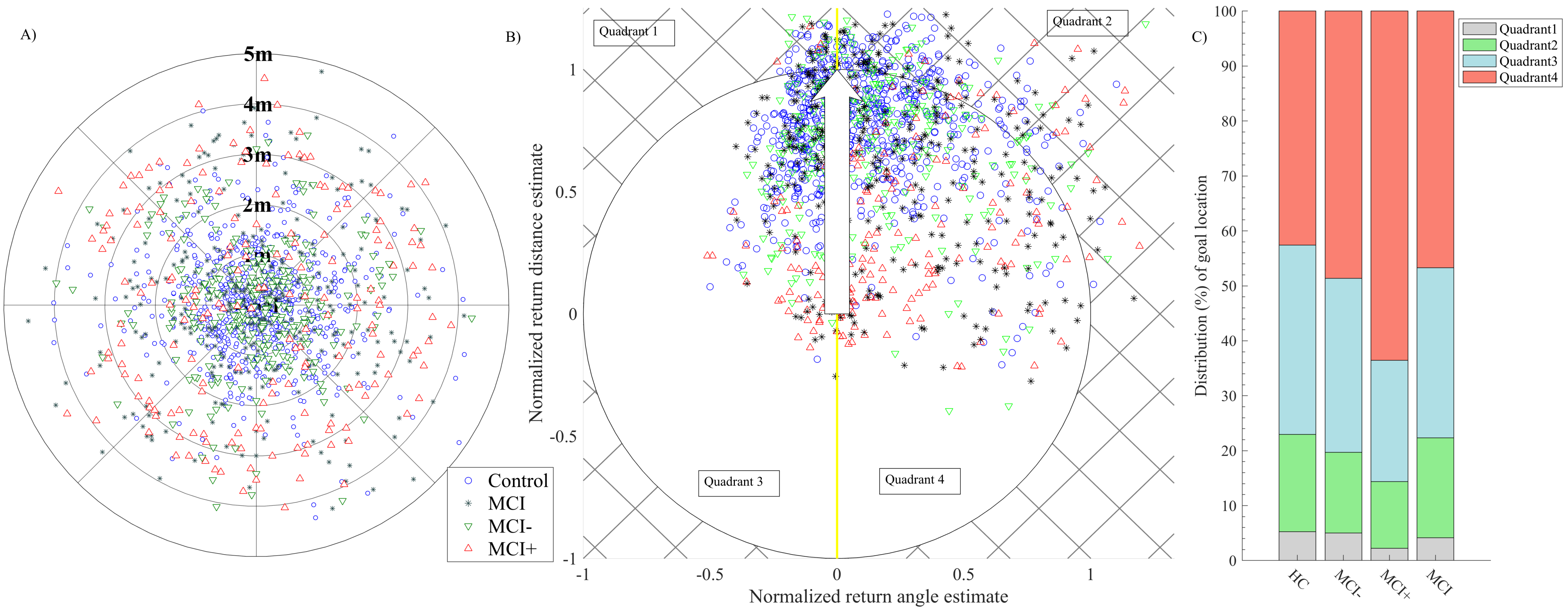


**Supplementary Fig 5. Distribution of goal location (cone 1) errors by group. A)** Illustration of absolute distance error from goal (cone 1, centre), each concentric circle represent 1 metre of absolute distance error. **B)** Distribution of normalized estimated location of cone1 translated and normalized to Cartesian space. Tip of the white arrow (0,1) represents the optimal linear return path to location of goal (cone 1). The border of the circle indicates optimal distance estimate, outside = overestimate (quadrants 1 and 2), inside = underestimate (quadrants 3 and 4). Yellow line indicates optimal rotation toward goal performed at cone 3, left of yellow line = under-rotation (quadrants 1 and 3), right = over-rotation (quadrants 2 and 4). **C)** Distribution of estimated location of cone 1 by quadrant, expressed as a percentage of the group's total responses. Each marker represents the outcome of a trial which did not reach 'out of bounds': blue circles = HCs; black asterisks = MCI without biomarkers; red triangles = MCI+; green inverted triangles = MCI-.
